# High resolution structure of a partially folded insulin aggregation intermediate

**DOI:** 10.1101/716845

**Authors:** Bhisma N Ratha, Rajiv K Kar, Jeffrey R Brender, Bankanidhi Sahoo, Sujan Kalita, Anirban Bhunia

## Abstract

Insulin has long served as a model for protein aggregation, both because of the importance of aggregation in insulin manufacture and because the structural biology of insulin has been extensively characterized. Despite intensive study, details about the initial triggers for aggregation have remained elusive at the molecular level. We show here that at acidic pH, the aggregation of insulin is likely initiated by a partially folded monomeric intermediate whose concentration is controlled by an off-pathway micellar species. High resolution structures of the partially folded intermediate show that it is coarsely similar to the initial monomeric structure but differs in subtle details – the A chain helices on the receptor interface are more disordered and the B chain helix moves away from C-terminal A chain helix. The result of these movements is the creation of a hydrophobic cavity in the center of the protein that may serve as nucleation site for oligomer formation. Knowledge of this transition may aid in the engineering of insulin variants that retain the favorable pharamacokinetic properties of monomeric insulin but are more resistant to aggregation.

## Introduction

The accumulation of misfolded proteins is a common pathological feature of a number of devastating neurodegenerative disorders, such as Alzheimer’s and Parkinson’s disease, and several metabolic diseases, such as type II diabetes. Formation of amyloid usually requires a protein passes through several intermediate forms prior to achieve a β-sheet rich fibrillar structure.^*1-3*^ According to the amyloid hypothesis the formation of amyloid progress through an initial nucleation phase followed by an explosive growth phase.^*4-6*^ The oligomers formed during the nucleation phase have long been implicated in disease progression and cytotoxicity, under both *in vitro* and *in vivo* conditions.^*7-12*^ Hence obtaining structural understanding of amyloid intermediates will help to devise inhibitors, which can either be used for treatment or prevention.

Besides the pathogenic amyloid proteins, insulin, a peptide hormone having therapeutic importance, also displays high propensity to form amyloid fibril. Because it is readily available and fibrillizes easily under certain conditions, insulin is often regarded as an excellent model for protein aggregation, especially for the subset of amyloidogenic proteins such as β-microglobulin^*13*^ and transthyretin^*14*^ which are initially folded but unfold during the aggregation process.^*15*^ Insulin is generally stable in its zinc-bound hexameric form. In the absence of zinc at physiological pH insulin is primarily present as a conformationally stable dimer, however it can aggregate at elevated temperature or during prolonged storage. At a pH below 2 in the absence of zinc, the dimer dissociates into the monomeric form.^*16, 17*^ The monomeric protein is more physiologically active than either the dimer or the hexameric form. Considerable effort has been therefore been devoted into engineering fast acting insulin analogs that favor the monomeric structure over other forms.

Unfortunately, insulin is more prone to aggregation as a monomer than as a hexamer. The full molecular trigger for aggregation is not completely understood, but it is known that low concentrations of urea^*18, 19*^ and alcohols^*20*^ stimulate amyloid formation by destabilizing the native structure. Based on these findings it is likely that insulin partially unfolds at contact with the air water interface^*21*^ to expose hydrophobic regions of the protein that trigger eventual aggregation.^*22*^ Owing to its commercial importance and vulnerability to amyloid fibrillation, substantial research has been directed at understanding the structures along the insulin aggregation pathway. While models of the amyloid fiber exist from cryo-electron microscopy (cryo-EM)^*23*^ and x-ray crystallography by a fragment based approach^*24*^ and substantial progress has also been made in understanding on a macroscale, high resolution structures of intermediates have been lacking.

To address this problem, we have followed the entire time course of aggregation using a series of spectroscopic techniques. Using a NOE restraints from solution NMR from timepoints identified by thioflavin T (ThT) fluorescence and other methods, we are able to show there is a sequential loss of helicity in the A chain helices at the insulin receptor interface and a movement of the B chain helix that leads to exposure of hydrophobic residues. This partially unfolded intermediate eventually leads to another anti-parallel β-sheet intermediate and may represent the first step along the insulin aggregation pathway.

## Results and Discussion

### Changes in Secondary Structure during Aggregation and Fiber Formation are Tightly Linked

The fibrillation kinetics of insulin can be followed directly by Thioflavin T, a well-established fluorescent probe for studying the kinetics of protein amyloid formation. Thioflavin T binds to the grooves formed between side chains of the β-sheet.^*25*^ The formation of the groove requires a regular lattice of side chains on the surface of the β-sheet. Because of this requirement, Thioflavin T is fairly specific for amyloid fibers, which have a regular linear structure, over other β-sheet morphologies. A more general measure of conformational change can be obtained from measurements of secondary structure from circular dichroism spectra. Since β-sheet structure is increased in most, but not all,^*26*^ aggregation intermediates, a comparison of the two methods provides an indication of whether non-amyloid intermediates are formed during aggregation. A close correspondence between the two methods is evidence, but not proof, that an appreciable population of non-amyloid oligomers is not formed during aggregation.^*27*^ Conversely, the Thioflavin T lagging behind the CD signal is strong evidence for a non-fiber β-sheet intermediate.^*28*^

Initially the CD spectra displays the two negative maxima at 222 nm and 208 nm (**Fig. 1A**) characteristic of the largely alpha helical conformation of the protein seen in the crystal structure. Deconvolution of the CD spectra using BeStSel^*29*^ confirms that the protein initially possesses a large proportion of alpha helix (**Fig. 1B**) and a very low proportion of β-sheet. At approximately 40 h, the helicity of the protein rapidly decreases in concert with an increase in β-sheet secondary structure until a mostly β-sheet structure is achieved at around 52 h (**Fig. 1B**). The thioflavin T fluorescence assay also shows the sigmoidal growth kinetics typical of amyloid proteins i.e. a quiescent lag period as nuclei are formed followed by rapid growth as fibers elongate and eventual saturation as the monomer pool is depleted (**Fig. 1C**). The kinetics can be fit to a Boltzmann Sigmoidal curve fitting equation with two kinetic parameters: a lag time (t_lag_) reflective of nucleation and a half-time (t_1/2_) reflective of the elongation rate (Equation 1).

**Figure 1.**
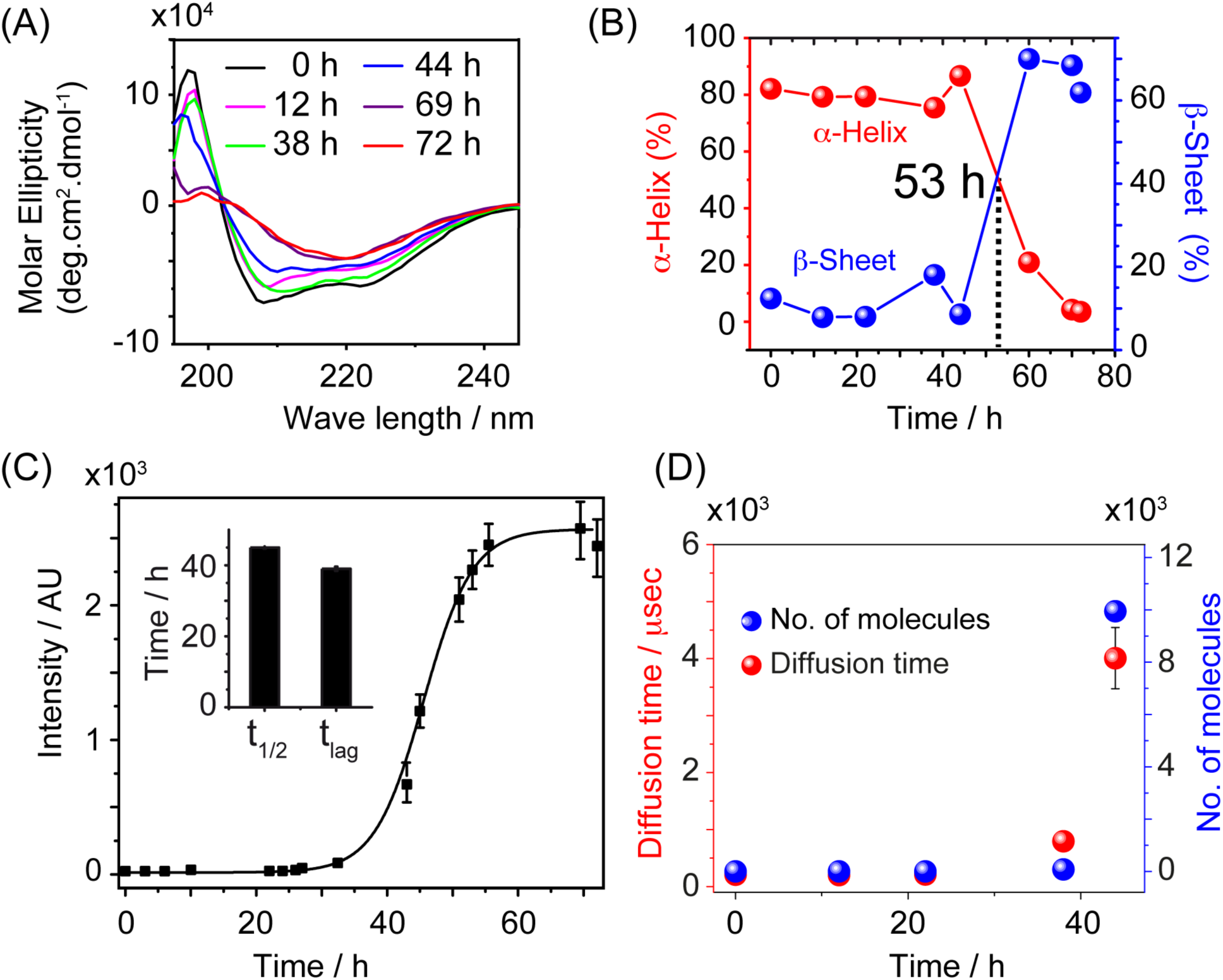
Amyloid formation, secondary structure changes, and protein aggregation happen on similar time scales. (A) Far UV CD spectra of 25 µM insulin in (aliquot from 2 mg/ml insulin in 20% acetic acid, Ph 1.9 at 335 K), diluted in 10 mM PBS pH 7.4, recorded at different time points. (B) BeSTSel deconvolution results of the CD spectra showing α-helix and β-sheet content plotted against the incubation time; the transition from α-helix to β-sheet time is around 53 h. (C) ThT fibrillation kinetics of 350 µM zinc free bovine insulin in 20% acetic acid (pH 1.9), inset shows the computed t_1/2_ (39 ± 0.6 h) and t_lag_ (45 ± 0.2 h) from Boltzmann fitting of the kinetics data. (D) Diffusion time and average number of molecules per particle derived from Fluorescence Correlation Spectroscopy (FCS) at different incubation times. For FCS, samples were incubated at 2 mg/ml (350 µM) and measured after dilution to 50 µM.

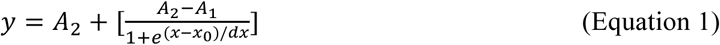

Both the calculated lag time of 39 ± 0.6 h and half-time (t_1/2_) of 45 ± 0.2 h approximately match rough estimates of the nucleation and elongation periods from the CD spectra. Overall, the results suggest a slow nucleation process and a relatively fast elongation rate in comparison to other amyloid forming proteins such as IAPP^*27*^ and Aβ.

The close correspondence between fiber formation suggests there is not a buildup of non-amyloid intermediates at any point during aggregation. However, the interpretation of the results is complicated by the difference in insulin concentrations used in each assay. To observe the aggregation progression directly in terms of growth in size of insulin amyloid aggregates, the diffusion time (τ_D_) of insulin specifically labeled with tetramethylrhodamine (TMR) at the N-terminus was measured by single molecule based fluorescence correlation spectroscopy (FCS) under similar conditions as the ThT assay. **Fig. 1D** represents the diffusion time (red dots) and its corresponding number of molecules in the aggregate (blue dots). A representative fitted curve with the fit residuals for each observed time point is displayed in **Fig. S1**. Fibrillation of insulin takes around 72 h for saturation; however, we were able to record the FCS spectra only up to 44 h, as after 44 h the suspension becomes too highly heterogeneous to determine the diffusion time. Although the heterogeneity limits detection at later time points, it can still be seen that the major increase of thioflavin fluorescence occurs nearly concurrently with the onset of aggregation in any form.

### Non-Fibrillar Oligomeric Intermediates Exist Early in the Aggregation Pathway

To investigate large aggregates not detectible by FCS, we employed time-lapse transmission electron microscopy (TEM) and atomic force microscopy (AFM) to visualize the progression of insulin fibrillation. As shown in **Fig. 2A**, no aggregates were detected in TEM micrographs during the lag phase of the CD and ThT assays (0 h and 18 h). Since the ThT assay is known to produce false positives under some conditions, we confirmed insulin was forming amyloid fibers under our conditions by examining the ultrastructural organization of the sample at the final time-point. As expected, both TEM (**Fig. 2A**) and AFM images (**Fig. 2B**) at 72 h shows a network like structure of straight fibers of ∼10 nm thickness, which is typical of mature amyloid insulin fibers.^*7,30,31*^ By contrast, AFM images at intermediate time points prior to entering the log phase of the aggregation show new intermediates distinct from either the initial monomeric protein or the mature amyloid fibers. At 38 h, spherical aggregates of ∼10 nm height are the dominant detectable species (**Fig. 2B**). Around the t_1/2_ (44 h) of the insulin aggregation, this intermediate was converted into proto-fibrillar structure consisting of relatively short ∼14 nm maximum heights was observed, similar to previous reports in the literature.^*23, 32*^

**Figure 2.**
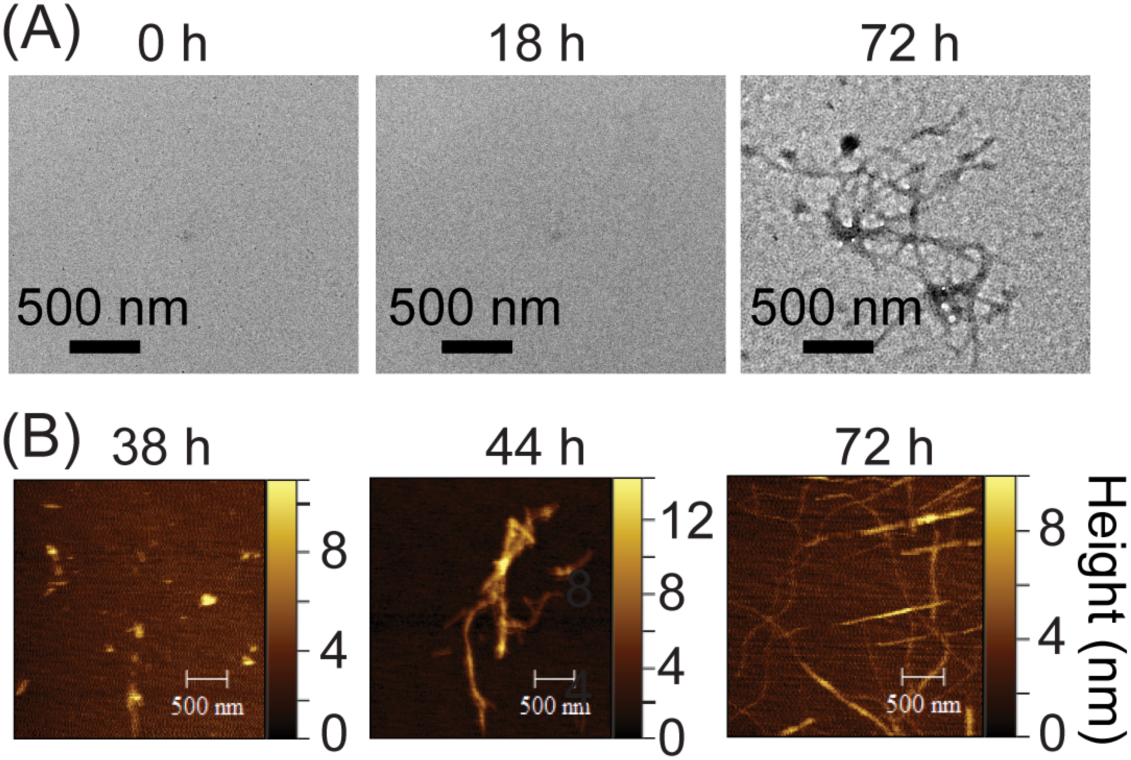
Non-fibrillar oligomeric intermediates in the insulin pathway. (A) TEM micrographs and (B) AFM images of 350 µM zinc free bovine insulin after incubation for the indicated time periods. While nothing can be detected in the lag phase (0 and 18 h), spherical intermediates of ∼10 nm in height are detectable before amyloid fibers form completely (72 h).

### Aggregation likely proceeds from a monomer derived species

The results of AFM and TEM can be highly dependent on both the type of surface used and the deposition method.^*33*^ Since oligomers not tightly bound to the substrate may detach from the surface during drying, measurements, microscopy measurements of oligomer formation may be biased by preferential binding. In particular, while micellar type intermediates have been identified as potential nucleation sites for amyloid formation,^*34-38*^ they may be missed by microscopy because the rapid dissociative equilibrium that characterizes a micelle often results in them being deposited as monomers on the mica surface.

To detect possible oligomers missed by TEM and AFM, we employed the classical fluorescent probe pyrene. Excitation of pyrene at 343 nm results in four emission peaks at 374 nm (I_1_), 378 nm (I_2_), 387 (I_3_) nm and 394 nm (I_4_). At low concentrations, the ratio of intensities of vibronic peaks I_1_ to I_3_ in pyrene is sensitive to the polarity of the local microenvironment with the I_1_/I_3_ ratio decreasing as the hydrophobicity of the environment increases, as would be expected when the highly hydrophobic pyrene binds to the hydrophobic interior of a disordered micelle.^*39*^ Similar to detergents, an apparent critical micellar concentration CMC can be defined by the inflection point in a plot of this ratio as a function of the protein concentration.^*40*^

**Fig. 3A** shows the fluorescence spectra 1 µM pyrene with increasing concentrations of zinc free bovine pancreatic insulin from 0 to 100 µM. Low concentrations of insulin below ∼ 10 µM have a I_1_:I_3_ ratio similar to that of pyrene in water (I_1_/I_3_ = 1.8). Above ∼ 50 µM, the I_1_:I_3_ ratio decreases substantially and becomes similar to that of pyrene in cyclohexane (I_1_/I_3_ ∼1.65).^*41*^ When the I_1_:I_3_ ratio is plotted against the concentration of insulin using Equation 2, an apparent CMC for zinc free insulin in 20% acetic acid of 45.82 µM was found (**Fig. 3B**). This value is higher than the known CMC values of other amyloid proteins, all of which undergo a transition from a largely disordered monomeric state to a partially folded conformation as a first step in the aggregation process. Insulin, by contrast, is initially folded and is less likely to self-associate into micellar forms. The existence of a CMC explains the flat concentration dependence observed in **Fig. 1**; the nearly similar apparent aggregation rates at 25 µM and 350 µM are consistent with aggregation proceeding from a monomer derived species whose concentration is held nearly constant by a micelle/monomer equilibrium.

**Figure 3.**
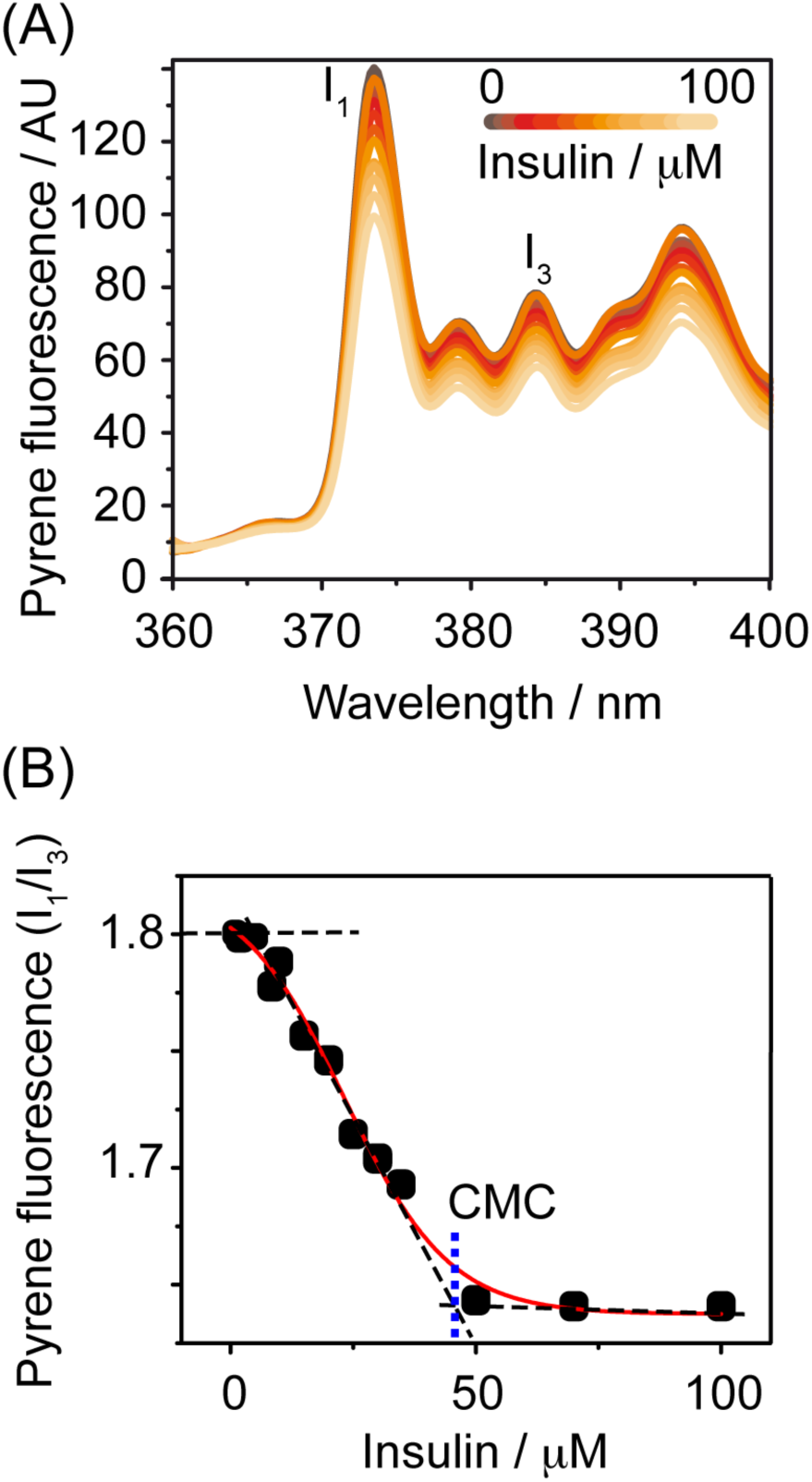
Insulin forms micelle-like oligomers at micromolar concentrations. (A) Emission spectra of 1 µM pyrene as a function of insulin concentration in 20% acetic acid. The environmentally sensitive I_1_ and I_3_ vibronic bands are indicated. The ratio of I_1_ to I_3_ decreases as the pyrene becomes bound in a hydrophobic environment. (B) Plot of the ratio of the I_1_ and I_3_ vibronic bands against the insulin concentration. The apparent CMC value (45.82 µM) determined by fitting to Equation 2 is indicated with the arrow.

### Monomeric Insulin Partially Unfolds During Aggregation

The previous results appear to suggest amyloid formation by insulin is a two-step process. However, the apparent mechanism can be sensitive to the method of detection. The thioflavin T assay is specific only to the cross-beta sheet structure of the fiber and will not detect any intermediates that lack this specific topology. Similarly, CD and FCS are insensitive to the presence of any intermediates that do not involve a change in secondary structure or size, respectively. The NMR chemical shift, by contrast, is highly sensitive to small changes in conformation,^*42*^ solvent exposure,^*43*^ electrostatics, and dynamics. NMR may therefore detect intermediates defined by subtle structural changes that are not obvious by other means, provided that they are small enough to be detected by NMR.^*44*^

The 1D ^1^H NMR for 2 mg/ml samples at 335 K is shown in **Fig. 4A**. Until the final time point at 73 h the spectra are strikingly similar, an indication that most of the population remains in a state that resembles the original monomeric protein. Two major changes are evident as the sample evolves. First, there is a 39% drop in overall intensity within the first 12 h after the start of aggregation before the system stabilizes (**Fig. 4B**). Second, a set of new peaks appears near 0 ppm, a region of the spectrum associated with mobile alkyl chains that are solvent protected and thus shifted to the high field region of the 1D ^1^H spectrum.^*45*^ This peak first appears 18 h after and is most prominent in the spectra at 1 h, gradually broadening and decreasing in intensity until it disappears completely at the 64 h point. Similar peaks have been detected in multiple other amyloid proteins where they have been associated by diffusion edited NMR spectroscopy with large spherical oligomers ∼50 nm in diameter.^*27, 46-49*^

**Figure 4.**
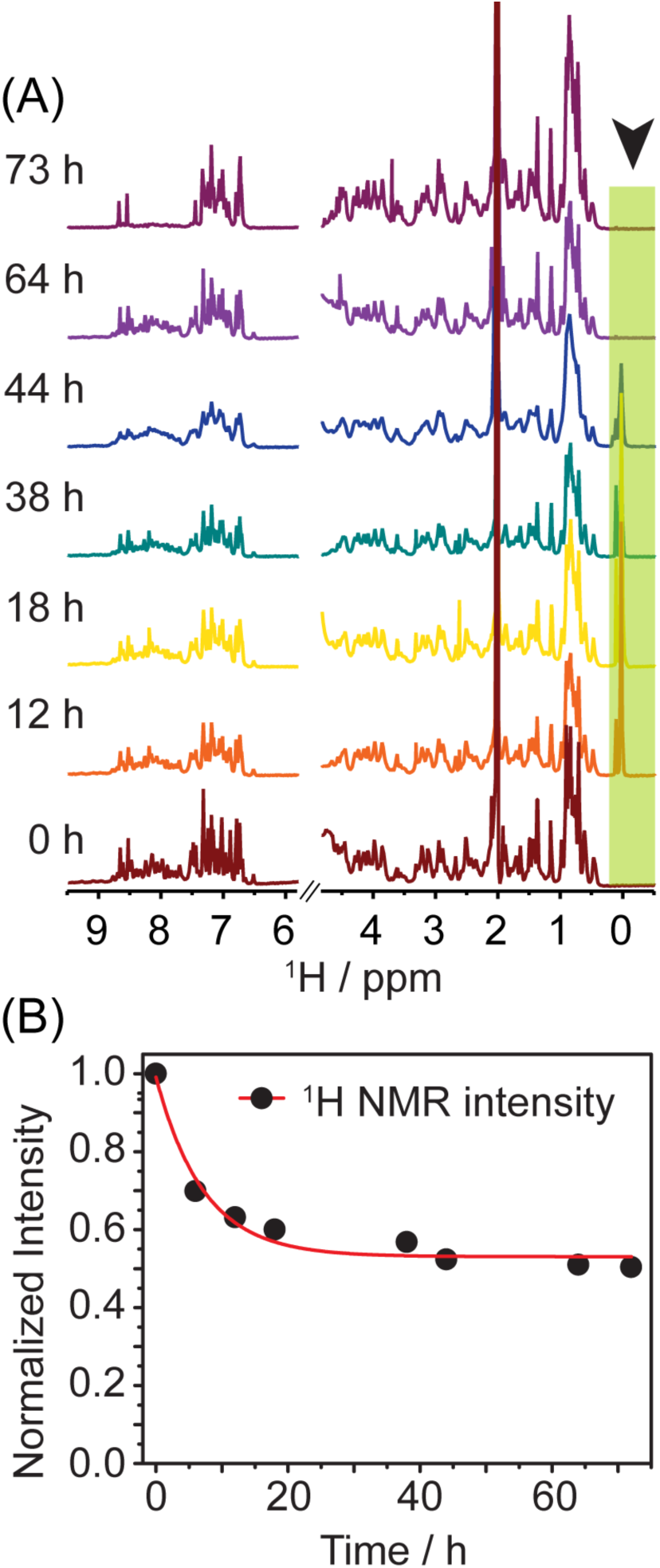
NMR shows an intermediate not detected by CD or ThT. (A) 1D ^1^H NMR spectrum as a function of time. A new peak appears near 0.5 ppm at 12 h that disappears completely by 64 h. (B) Intensity of the aromatic and amide region of the proton NMR spectrum showing the gradual loss of NMR observable species with increasing incubation time. The experiment was performed using Bruker AVANCE III 700 MHz, equipped with a cryoprobe.

The chemical shift of Hα proton gives valuable information on the local chemical environment of the residue when compared to its known values in a random coil.^*50, 51*^ To measure fluctuations in the local chemical environment of the monomeric protein, we compared the Hα chemical shifts at a series of time points to the starting spectra (t=0) (**Fig. S2**). The chemical shift changes are small and in almost all cases below the 0.1 ppm threshold used to identify secondary structure. This suggests the changes primarily involve relatively modest rearrangements of existing structural elements rather than large scale movements or changes in secondary structure. Within the structure, the changes in chemical shift are most evident at later time points and are primarily localized to the B chain alpha helix from H10 to L15 (**Fig. S2**).

To resolve fine structural details, a series of 2D NOESY spectra were taken at each time point (**Figs. 5A-B**). The intensity of NOESY cross-peaks is sensitive to conformational exchange and provides complementary information to the chemical shifts. The alpha helical signature NOEs such as αN (i to i+3/i+4) and αβ (i to i+3), corresponding to the crystal structure of insulin are particularly informative in this regard. In particular, the L13^A^α/L16^A^β peak gradually broadened with time and the NOE completely disappeared after 44 h of incubation, around the t_1/2_ of the ThT kinetics. Similarly, the L16^A^α/Y19^A^β, V12^B^α/L15^B^β, and H10^B^α/A14^B^H peaks steadily broaden with time. Overall, the ends of the B chain helix show a time dependent decrease in crosspeak intensity and change in chemical shifts that suggest a gradual loss of secondary structure at the ends of the helix (**Figs. 5C-E**).

**Figure 5.**
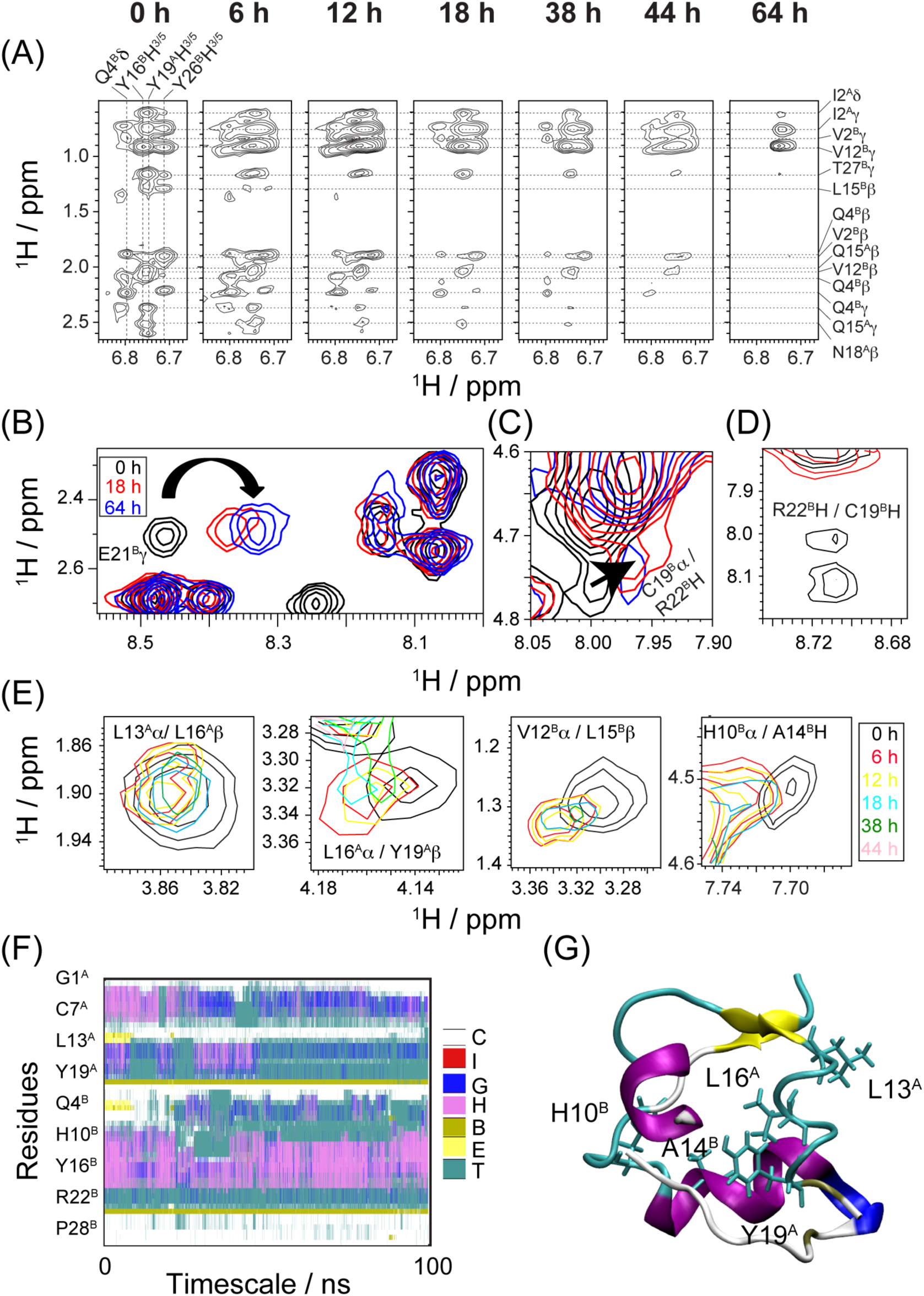
Sequential unfolding of insulin with increasing incubation time. (A) Aromatic ring region of the 2D NOESY spectrum showing the broadening of inter-chain and long-range intra chain NOEs as incubation time increases. (B) Close up of the peak for E21^B^γ located on the loop connecting the B chain central helix with the C-terminal with color coded time points. (C) NOE between C19^B^α proton and amide proton of R22^B^ showing chemical shift perturbation and gradual broadening with time. (D) NOE between C19^B^H/R22^B^H showing chemical shift perturbation and gradual broadening with time. (E) Alterations in the α-helices as time progresses shown by chemical shift perturbations and gradual broadening of α-helix signature NOEs color. (F) Changes in the molecular conformation (secondary structure) of the initial starting structure as a function of time from MD simulation. (G) Representative snapshot from trajectory, displaying important side chain conformations.

Long range NOEs (i to i+ ≥5) and side chain NOEs provide essential information on the tertiary structure of the protein. At the initial time point (0 h), the NMR structure is defined by both inter-chain NOEs (such as I2^A^δ or I2^A^γ/Y26^B^H^3/5^ and L15^B^β/Y19^A^H^3/5^) and long range intra A chain NOEs including L15^B^β/Y26^B^H^3/5^, I2^A^δ or I2^A^γ/Y19^A^H^3/5^, V12^B^/F24^B^, L15^B^/F25^B^, G8^B^/Y26^B^ and V12^B^/F24^B^ (**Table S1**). The gradual broadening of these NOEs are shown in the subsequent panels of **Fig. 5A**. Interestingly, there are no NOE cross-peaks associated with the intense peak at 0 ppm that appears and disappears as aggregation progresses. This suggests the peak arises from a separate species that is independent of the insulin monomer, as would be expected if it was associated with an oligomeric intermediate.

As the NMR results suggest a critical event in the aggregation of insulin is a partial unfolding of the insulin monomer, we solved the structure at 0, 18, and 38 h using the NOE and torsion angle restraints available at each time point (**Table S2**). The structure at the initial time point (PDB 6KH8) still closely resembles the T state of the insulin hexamer (**Fig. 6**),^*52*^ in agreement with previous studies of the insulin monomer.^*53, 54*^ The A chain is comprised of two helices from I2^A^ to T8^A^ and L13^A^ to C20^A^ connected by relatively rigid linker region constrained by an intra-chain disulfide bond in an antiparallel helix-loop-helix motif. The most prominent feature of the B chain structure is central helix from S9^B^ to V18^B^. The helix in B chain is flanked by two inter-chain disulfide bridges with the remainder of the molecule in an extended conformation. The helices of the A chain are positioned nearly perpendicular to the B helix. In combination with the B chain C-terminal strand, this arrangement creates a small hydrophobic core at the A/B chain interface that is at least partially shielded from solvent, although the inter-chain disulfide bonds are essential for the stability of the protein.

**Figure 6.**
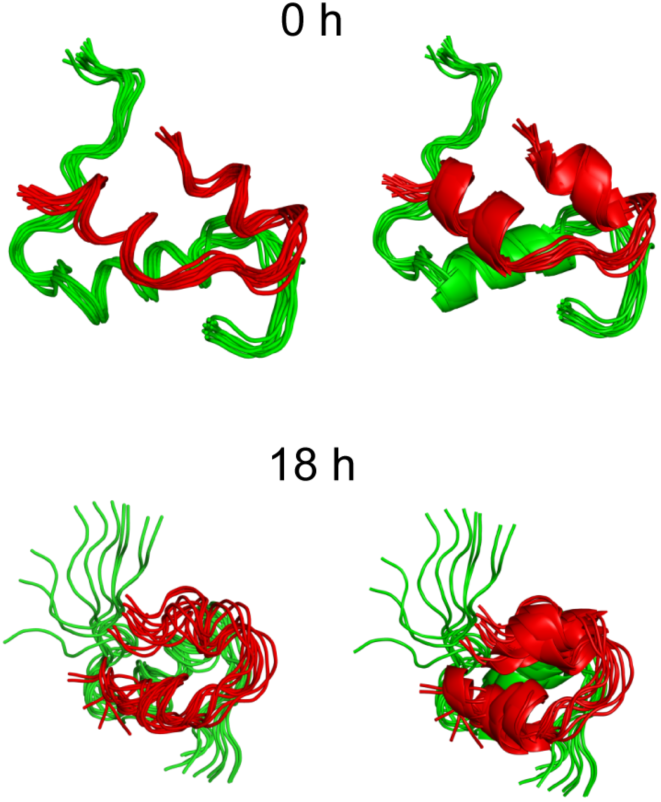
Three-dimensional solution structures of bovine insulin at 0 and 18 h. The structural ensembles were created through 100 ns molecular dynamics using the ff99SBuildin force-field and NOE constraints from NOESY spectra at the indicated time points as restraints.

Starting from the same initial structure but using NOE restraints from different time points in the aggregation experiment, it is possible to see the partial unfolding of the monomer as time progresses. When the NOE restraints from the initial 0 h time-point are used, a network of contacts between hydrophobic residues (I2^A^ and Y19^A^ and Y26^B^, L15^B^ and Y19^A^, and V12^B^/F24^B^) defines the tertiary structure (green residues in **Fig. 7A**). These contacts are partially lost when restraints from the 18 h time-point are used (PDB acquisition code 6KH9). While the contacts between the δ/γ protons of I2^A^ and H^3/5^ of Y19^A^ were unchanged, the distance between the other contacts within this hydrophobic cluster increases (green residues in **Fig. 7B**).

**Figure 7.**
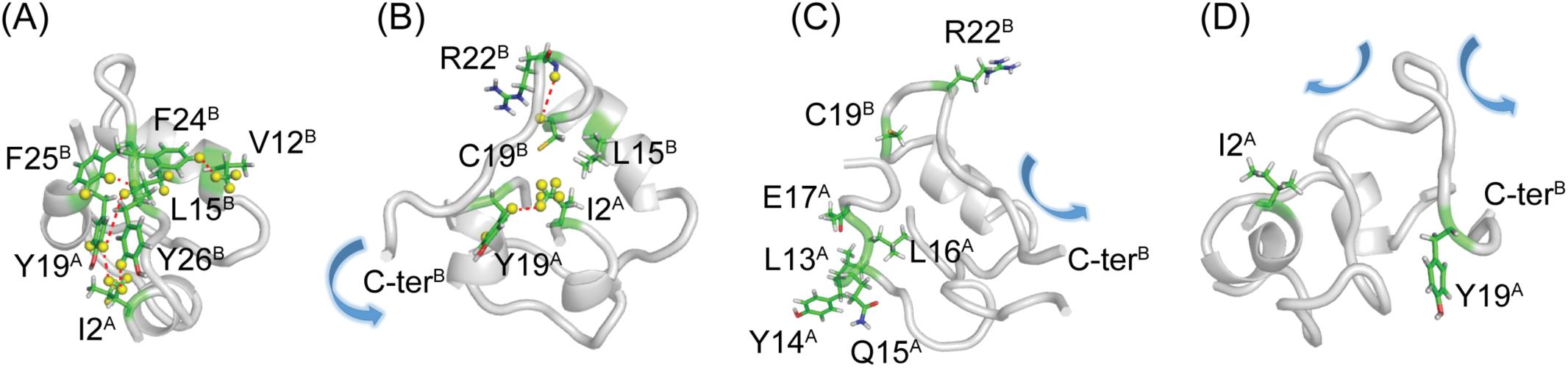
Three-dimensional structures at different time points. **(A)** The initial structure at 0 h (PDB: 6KH8), (B) 18 h (PDB: 6KH9), (C) a high temperature intermediate (PDB: 1SF1) and (D) 38 h (PDB: 6KHA). Important inter-residue proton contacts are shown with dashed lines (yellow spheres). Movements of the main chain relative to the monomeric structure in chain dynamicity are shown with blue arrows.

The consequences of the loss of these contacts is an opening up of the structure, creating a hydrophobic cleft in the center of the protein bounded by F24^B^, F25^B^, Y26^A^, V12^B^, V3^A^, and Y19^A^ (**Fig. 8**). One major difference between the two structures is a rotation of the C-terminal end of the B chain helix away from the C-terminal A helix with the loss of the hydrophobic contacts between L15^B^ and V19^B^ on the B chain helix and L16^A^ on the C-terminal A helix.

**Figure 8.**
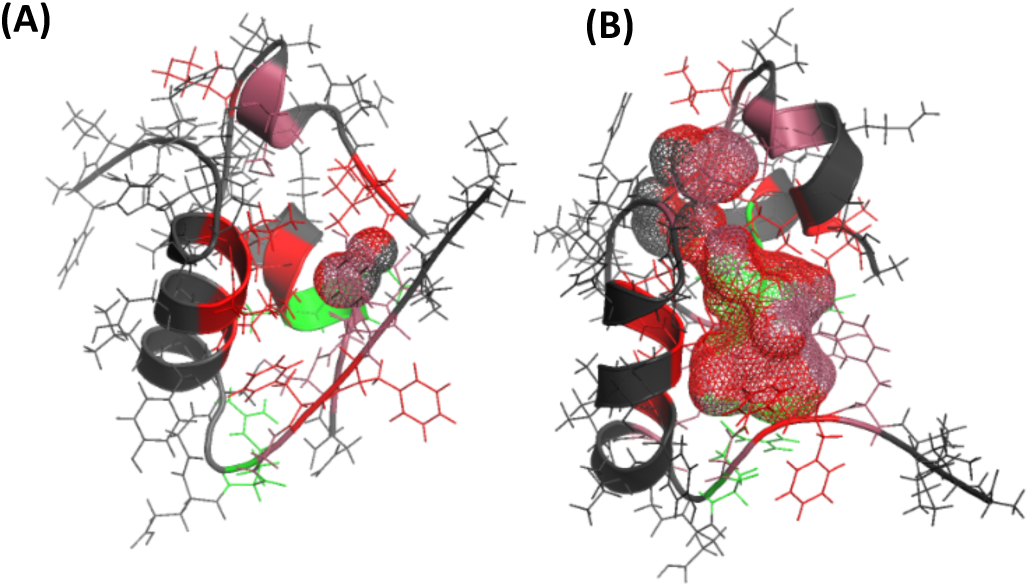
Movement of the B chain relative to the C-terminal A chain helix creates a hydrophobic cavity. Residues are colored by approximate hydrophobicity scale – red hydrophobic (V,I,L,F), raspberry neutral (Y,G,C), green polar or charged (Q,R,N). Residues outside the center are in black. (A) Representative structure at the start of the incubation period. The structure is well folded with hydrophobic residues (red) packed tightly against each other. (B) Representative structure after 18 h of incubation. A large cavity lined by hydrophobic residues exists in the center of the protein.

This change occurs simultaneously with a partial loss of helicity on the C-terminal A helix, which begins to unravel as the A chain adopts a turn conformation at the C-terminus. This unraveling is partly explained by the loss of some contacts between the C-terminal strand of B chain and the A chain helices as the C-terminal end of the strand bends away from the A chain helices and towards the newly positioned B-helix. The movement is confirmed by differences in chemical shifts in the loop connecting the B chain C-terminal strand and helices (**Fig. 5B**). This motion, a hinge-like rotation about a conserved β-turn from G20^B^ to G23^B^, is somewhat analogous to the motion insulin undergoes in binding to its receptor.^*55*^ Interestingly, while the unwinding of the A chain helices is found in a high temperature structure of insulin (PDB 1SF1, **Fig. 7C**), the disengagement of the C-terminal strand of B chain and rotation of the B chain helix are not, suggesting these movements are more important for rapid aggregation than A chain helix unwinding.^*22*^

While NMR can give information about the structure of the partially unfolded intermediate, it cannot resolve the sequence of motions that give rise to it. To gain a greater understanding of this process, we used an all-atom MD simulation of the initial insulin monomer structure starting from three independent trajectories with similar initial atomic coordinates but varying initial velocities without using experimental NOE restraints. The simulation was performed at low pH to correlate with the experimental condition.

The monomer structure is relatively stable overall, as shown by a plot of the secondary structure over the course of the simulation. In particular, the B chain helix remains a helix throughout the full 100 ns of simulation (**Fig. 5F**). The A chain helices, by contrast, begin to unwind at around the 50 ns mark (**Fig. 5G**). Conformational analysis from simulation also reveals high fluctuations for the C-terminal residues of B chain. Both motions are indicative of a model where the partial unfolding of the insulin monomer is initiated by the movement of the B chain C-terminal strand away from the A chain helices, followed by the destabilization of the A chain helices after the loss of contacts with the B chain C-terminal strand.

### A late stage intermediate contains an anti-parallel β-sheet

The structures at later time points are poorly defined as most of the NOE restraints have been lost (PDB 6KHA). The B chain helix has begun to unfold and what little tertiary structure exists is primarily by the constraints introduced by the three-disulfide bonds (**Fig. 7D**). The conformational variability at later time points was confirmed by deuterium exchange NMR experiments (**Fig. S3**). In contrast to the initial time 0 sample, where peaks corresponding to B chain helix can still be detected even 6 h after exchange into 80% D_2_O, the sample incubated for 64 h exchanges its protons relatively rapidly, indicating intact B chain helices are less prevalent in the population.

While a high-resolution structure is difficult to obtain, we attempted to identify the stretch of peptide responsible for adding the monomer to fibril mass, assuming that a monomeric or oligomeric species may have a conformation similar to that of the units of the fibril. We first searched the later time points for the appearance of β-sheet specific NOEs that have been previously identified in the literature. A few reports suggest the presence of parallel β-sheets^*56,57*^ in the insulin amyloid fibril, however, others suggest the presence of antiparallel β sheets.^*24, 58-60*^ Using a fragment-based approach, Ivanova et al. identified the helical region (L11^B^-V18^B^ in bovine insulin) of B chain as the minimal fragment of the insulin to form amyloid fibers and built a model for the amyloid fiber of the whole protein based on the crystal structure of the L11^B^-L17^B^ fragment.^*24*^ However, we were unable to find NOEs reflective of either parallel or anti-parallel β-sheets at any timepoint.

The slow tumbling of large oligomers results in a broad weak signal that might obscure potential β-sheet NOEs. To better detect NOEs due to β-sheet formation, we incubated the sample for 72 h in 80% D_2_O and 20% D_4_ acetic acid to form amyloid fibers and then carried out a NOESY experiment; this experiment helped to reduce the noise introduced by water in the NOESY spectra. We were unable to observe NOEs between amide protons, as the labile proteins likely exchanged during the incubation period prior to forming the beta sheet. Although we could not detect any amide protons in the sample, we did detect two new NOEs between the alpha protons of H10^B^α and V18^B^α and between V12^B^α and Y16^B^α (**Fig. S4**). It is unlikely that these peaks arise from direct observation of the amyloid fiber due to its very large size and long rotational correlation time. Instead, they likely arise from smaller β-sheet containing oligomers, possibly catalyzed by interaction with the fiber surface.^*61-63*^

While we could not make a definitive assignment due to the lack of i,i+2 NOEs, it is possible to check the consistency of the two new NOE peaks with the model of the insulin amyloid fiber proposed by Ivanova et al. Both H10^B^ and V18^B^ and V12^B^ and Y16^B^ are in close in contact with each other in the model in which the the helical region of B chain forms anti-parallel β-sheets along the fiber axis (**Fig. S4**). Although our other models can be made that are also consistent with the data, the NOE peaks do lead credence to an anti-parallel β-sheet species being present.

## Conclusion

Protein fibrillation is a multistep process that progresses through a series of transient structural intermediates. Mostly the transient oligomeric forms of aggregates are toxic. Lack of sufficient structural details of these toxic oligomers hinders effective drug designing and treatment approach. We have used zinc free insulin in this study to decipher the structural intermediates of insulin amyloid formation. As shown schematically in **Fig. 9**, insulin in strongly acidic buffer conditions (20% acetic acid-d4, pH 1.9) initially assumes a well-folded, compact, monomeric conformation (**Fig. 6**) that progresses through series of structural transitions on the path to becoming an amyloid fiber. At concentrations above ∼40 µM (**Fig. 3**), the monomer is in fast equilibrium with a micelle like species off pathway form the main amyloid formation route that serves to buffer the concentration of the aggregation prone monomer (**Fig. 9B**). The monomer itself is conformationally unstable in this condition, and the well-folded state coexists with partially unfolded conformations (**Fig. 9C**). The difference between the partially unfolded state and the native conformation is relatively subtle in comparison to later stages of aggregation (**Fig. 6**). The partially unfolded intermediate has a close enough secondary structure to the native protein that its appearance is not obvious by circular dichroism, giving the appearance of a simple two step transition (**Fig. 1**). While the differences between the two structures are relatively minor, the consequences are substantial. The movement of the B chain C-terminal strand away from the main body of the protein along with a rotation of the B chain helix (**Fig. 7**) creates a hydrophobic cavity in the protein that is not present in the native structure (**Fig. 8**), producing a potential binding site for aggregation. Aggregates (**Fig. 9D**) are detectable as a distinct peak in the NMR spectrum (**Fig. 4A**) uncoupled with any others at around 12 h after being dissolved into solution that is correlated with the transient appearance of spherical oligomers in AFM images (**Fig. 2B**). As time progresses, this large oligomer disappears and a new oligomer characterized by anti-parallel β-sheet formation emerges (**Fig. 9E**). By identifying and characterizing structural intermediates in insulin amyloid formation, this study will hopefully assist in understanding similar events of amyloid fibrillation in other amyloidogenic protein sequences and in the structure based design of robust insulin amyloid inhibitors.

**Figure 9.**
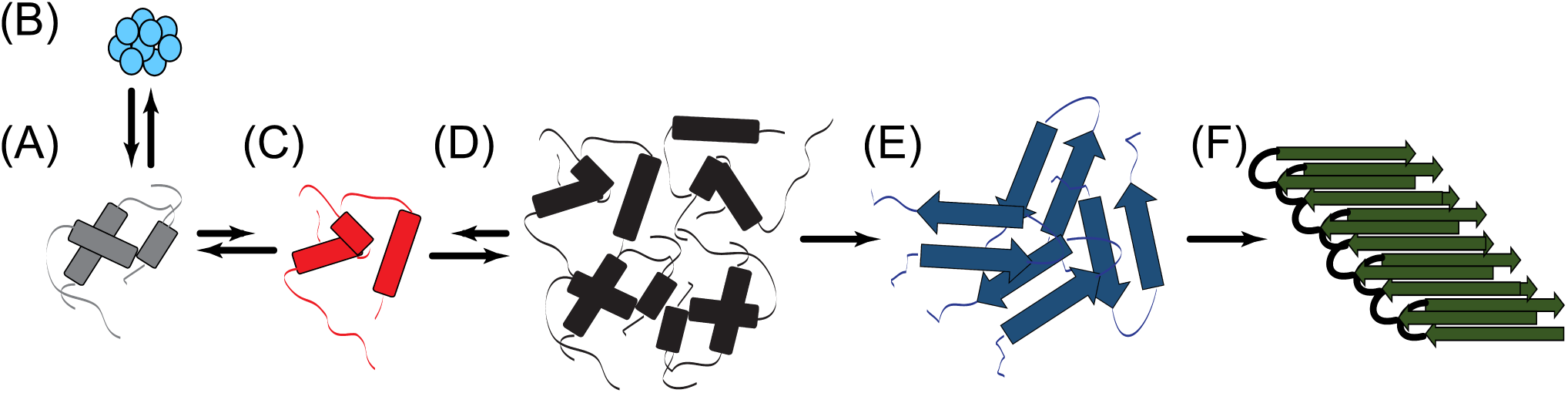
Simplified schematic of the proposed structural transitions on the path to amyloid formation. The monomeric protein (A) exists in fast equilibrium with insulin micelles (B) at high concentration and a partially unfolded state (C). The partially unfolded state is stabilized by binding to other insulin molecules and eventually forms a larger oligomer (D) which is sequentially converted into an anti-parallel β-sheet containing oligomer (E) and finally the amyloid fiber (F).

### Experimental procedures

Bovine pancreatic insulin was procured from Sigma (Missouri, USA) (Catalogue no. I5550). Tetradutero acetic acid (d_4_-acetic acid) and deuterium oxide (D_2_O) was purchased from Cambridge Isotope laboratories (Massachusetts, USA). TMR-succinimidyl ester was purchased from Invitrogen (California, USA). Reagents used in all other experiments were at least of ACS grade purchased either from Sigma (Missouri, USA) or Merck (New Jersey, USA).

### TMR labeling of Insulin

Bovine insulin was made zinc free as described earlier.^*31*^ 1 mg of protein was dissolved in 10 mM phosphate buffer pH 7.4 at a final concentration of 2 mg/ml. The resultant solution was degassed to avoid air oxidation of dye. To the protein solution, an aliquot of TMR succinimidyl ester was added slowly at room temperature with stirring. The protein-dye mixture was allowed to react overnight at 4 °C. After completion of the reaction the protein was separated from unreacted dye by size exclusion chromatography using Superdex Peptide 10/300 GL (GE Healthcare, Illinois, USA). The aliquots were collected, and concentration of labelled protein was determined. Then the aliquots were flash frozen in liquid nitrogen and stored at −80 °C freezer until further use.

### ThT Fluorescence Assay

Bovine pancreatic insulin was solubilized in 20% Acetic Acid at a final concentration of 2 mg/ml and pH adjusted to 1.9 with HCl. The UV absorbance determined the concentration of insulin in solution at 276 nm; bovine insulin has an extinction coefficient of 0.91 mg/ml for 1 OD at 276 nm. ThT was dissolved in miliQ water and centrifuged at high speed to get rid of the undissolved ThT. The concentration of ThT solution was determined by using a molar extinction coefficient of 36,000 M^−1^ cm^−1^. 2 mg/ml insulin solution was incubated at 335 K in a water bath; aliquots were drawn at different time points. The aliquots were diluted in 10 mM phosphate buffer (pH 7.4) with 100 mM NaCl.^*16, 64*^ The ThT fluorescence was measured in a quartz cuvette; the spectrofluorometer (PTI) was set at an excitation wavelength of 450 nm, and the emission was observed at 482 nm.

Observed fluorescence intensity at 482 nm for different time points was plotted against respective time points, and the curve was fitted by Boltzmann equation (Equation 1),

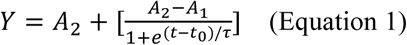

where, A_1_ is the initial fluorescence, A_2_ is the maximum fluorescence, t_0_ stands for the time where the fluorescence has reached to half of the maximum value, and 1/τ is the apparent rate constant of fibril growth and lag time approximated to t_0_-2τ.

### Pyrene CMC Assay

1 M pyrene solution in DMSO was prepared and concentration was confirmed by the help of UV absorption spectroscopy with a molar extinction coefficient of 54000 M^−1^ × cm^−1^. Pyrene was diluted to 1 µM in 20% acetic acid, and desired concentration of protein was added to it. Fluorescence spectra of free pyrene and in the presence of variable concentration of protein was recorded at an excitation wavelength of 343 nm. The emission spectra were recorded the I_1_/I_3_ ratio was plotted against the protein concentration and further analyzed by Boltzmann function as shown in equation 1 to determine the CMC value.

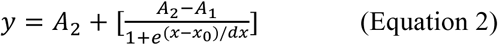

Here y corresponds to the ratio of the pyrene I_1_ and I_3_ vibronic bands, x is the concentration of the protein, A1 and A2 are the upper and lower limits of the sigmoid, x_0_ is the center of the sigmoid and dx is directly related to x in the region of abrupt change. The CMC is defined as the intercept of the tangent to the curve at x_0_.

### FCS Experiment

The 2 mg/ml fibrillation mixture of zinc free bovine pancreatic insulin doped with 1% TMR labeled insulin. The sample was diluted to 50 µM in the same fibrillation buffer i.e. 20% acetic acid, pH 1.9. 100 µl of the sample were taken in glass chamber slide and, was excited with a 532 nm laser on a vertical FCS setup at room temperature. The sample was allowed to settle for few minutes before recording the spectra to avoid contribution sample turbulence. The data obtained were analyzed as per the following equation and scans fitting with Chi-square > 0.999 were chosen to prepare the report as these spectrums were representing the major proportion of the sample.

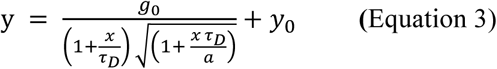

Here, G(τ) is autocorrelation function, G(0) is the inverse of the number of diffusing molecules in the confocal volume, “a” is constant dependent on the ratio of the radial and axial dimensions of the confocal volume, and τ_D_ is the diffusion time of the molecules.

### Circular Dichroism (CD) Spectroscopy

During insulin amyloid fibrillation the changes in the secondary structure of insulin were observed with the help of CD spectroscopy. The far-UV CD spectra were recorded on a quartz cuvette (0.2 cm) with the aid of Jasco J-815 spectrophotometer. 10 mM PBS (pH 7.4) was used throughout the study to adjust the concentration and spectrophotometer cell temperature was set at 298 K. The CD spectra recorded over a range of 200-260 nm with 1 nm data interval and scanning speed of 100 nm/min. Each CD spectrum represents an accumulation of four subsequent scans. The recorded raw spectra data in millidegrees were subtracted from the blank buffer and transformed to molar ellipticity using the following equation (Equation 4).

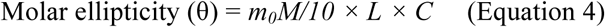

Here, m_0_ represents millidegrees, M stands for molecular weight (g mol^−1^), L is the path length of quartz cuvette used (cm), and C is the molar concentration of protein.

### NMR Experiments

The NMR experiments were performed at 298 K on Bruker AVANCE III 700 MHz spectrophotometer equipped with QCI cryoprobe. Each experiment was performed on a separate, independent sample incubated for the time indicated. For data acquisition and processing Topspin v3.1 was utilized. For all the experiments Zn-free insulin was used. 1D ^1^H experiments were recorded with two-fold serial dilution and accordingly a two fold increase in the number of the scan were made for the subsequent samples. The spectrum were normalized based on the following equation.

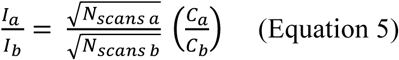

where, I_a_ and I_b_ stands for the intensity of the 1D ^1^H NMR spectra, N stands for the number of scans for samples a and b. C stands for the molar concentration of the sample a and b.

The sample were then dissolved in 20% tetradeutero acetic acid, 70% water and 10% D_2_O, pH 1.9 or 20% tetradeutero acetic acid and 80% D_2_O, pD 1.9 at a final concentration of 2 mg/ml and incubated for indicated time at 335 K, spectra were recorded immediately after completion of incubation in Shigemi tube. 2D ^1^H-^1^H homonuclear total correlation spectroscopy (TOCSY) and nuclear Overhauser effect spectroscopy (NOESY) were recorded at 298 K with 40 ms and 200 ms mixing time, respectively. A fresh sample was used for recording NOESY spectra of indicated time point. NOESY experimental parameters were kept the same as described earlier.^*31*^ The insulin TOCSY and NOESY spectra were analyzed with Sparky software (https://www.cgl.ucsf.edu/home/sparky).

### Molecular Modeling

3D structures of bovine insulin at different incubation time period were calculated using the NMR-derived MD simulation with distance and torsional restraints (obtained from the PREDITOR server from CαH chemical shifts), which were used for the calculation of the initial structure using the ff99SBuildin force-field^*65*^ in Amber 14 (**Table S2**). The starting coordinates for bovine insulin were obtained from X-ray structure of the insulin hexamer derived at 2.56 Å resolution (accession code: 2ZP6). Glu and His residues in the structure were protonated to correlate with the low pH conditions of experiment. The structure was first minimized using simulated annealing in vacuum for 20 ps with generalized Born model with the NMR derived restraints. The force constant values for the lower and upper bound distance and torsional restraints were varied from 10–32 kcal.mol.Å^2^. The temperature of the system was regulated using Berendsen temperature coupling algorithm^*66*^ and a cutoff of 15.0 Å is used for non-bonded interactions. The ensemble of the ten lowest energy conformations of insulin structure was used to represent the average structure of the protein. To test the stability and possible dynamics of the structure, three MD trajectories were collected for a time scale of 100 ns with similar initial condition but varying initial velocities. The SHAKE algorithm^*67*^ was used for re-ordering the hydrogen bond length during simulation with an integration time step of 2 fs The conformational analysis from the simulations were carried out using VMD.^*68*^

## Acknowledgments

This work was partly supported by Council of Scientific and Industrial Research (02(0292)/17/EMR-II to AB) and partly by Department of Biotechnology (BT/PR29978/MED/30/2037/2018 to AB) Govt. of India. BNR thanks UGC, Govt. of India for providing Fellowship.

## Notes

The three-dimensional bovine insulin structures at different incubation time have been submitted to Protein Data Bank (PDB ID: 6KH8 for initial time point (t=0 h); PDB ID: 6KH9 after incubation of 18 h; and PDB ID: 6KHA for insulin after incubation of 38 h).

## Author contributions

AB designed the research; BNR performed all biophysical experiments, analyzed the data with AB and did the assignment of all NOESY spectra; RKK performed MD simulation to determine the three-dimensional structure; JRB analyzed the data and edited the manuscript; BS and BNR performed AFM and FCS experiments; SK recorded TEM images; BNR, JRB, and AB wrote the manuscript. All authors reviewed the manuscript; AB arranged funding for this work.

## Abbreviations

cryo-EM: Cryo-electron microscopy
ThT: Thioflavin T
CD: Circular Dichroism
h: hour/s
t_lag_: Lag time
t_1/2_: Half-time
τ_D_: diffusion time
FCS: Fluorescence correlation spectroscopy
TMR: Tetramethylrhodamine
TEM: Transmission electron microscopy
AFM: Atomic force microscopy
CMC: Critical micellar concentration
NMR: Nuclear magnetic resonance
TOCSY: Total correlation spectroscopy
NOESY: Nuclear overhauser effect spectroscopy
MD: Molecular dynamics
d_4_-acetic acid: Tetradutero Acetic acid

